# The connectome of the *Caenorhabditis elegans* pharynx

**DOI:** 10.1101/868513

**Authors:** Steven J. Cook, Charles M. Crouse, Eviatar Yemini, David H. Hall, Scott W. Emmons, Oliver Hobert

## Abstract

Detailed anatomical maps of individual organs and entire animals have served as invaluable entry points for ensuing dissection of their evolution, development, and function. The pharynx of the nematode *Caenorhabditis elegans* is a simple neuromuscular organ with a self-contained, autonomously acting nervous system, composed of 20 neurons that fall into 14 anatomically distinct types. Using serial EM reconstruction, we re-evaluate here the connectome of the pharyngeal nervous system, providing a novel and more detailed view of its structure and predicted function. Contrasting the previous classification of pharyngeal neurons into distinct inter- and motorneuron classes, we provide evidence that most pharyngeal neurons are also likely sensory neurons and most, if not all, pharyngeal neurons also classify as motorneurons. Together with the extensive cross-connectivity among pharyngeal neurons, which is more widespread than previously realized, the sensory-motor characteristics of most neurons define a shallow network architecture of the pharyngeal connectome. Network analysis reveals that the patterns of neuronal connections are organized into putative computational modules that reflect the known functional domains of the pharynx. Compared to the somatic nervous system, pharyngeal neurons both physically associate with a larger fraction of their neighbors and create synapses with a greater proportion of their neighbors. We speculate that the overall architecture of the pharyngeal nervous system may be reminiscent of the architecture of ancestral, primitive nervous systems.

## INTRODUCTION

A detailed understanding of nervous system function, development and evolution necessitates precise anatomical descriptions of the individual components of a nervous system and how these individual components interact with each other on the level of individual cells and synapses. The cellular complexity of vertebrate brains makes such anatomical descriptions exceptionally difficult^1^, but even much simpler invertebrates contain nervous systems of astounding cellular complexity. This complexity problem is mitigated by focusing such anatomical and functional descriptions on individual units of a nervous system that display autonomous functions that can be studied in relative isolation. The stomatogastric ganglia (STG) of lobsters and crabs are prime examples for how the study of a simple functional unit of a nervous system can reveal fundamental insights into the generation of behavior^2^. However, many of these simple, self-contained circuits have been studied in animals that cannot be easily subjected to genetic or developmental analysis. The nervous system of the nematode *C. elegans* does not share these limitations. Its entire nervous system is composed of little more than 300 neurons^3^ that assemble into a complex, sex-specific neuronal wiring pattern^4,5^, but in analogy to the STG of crabs and lobsters, *C. elegans* also contains a simpler functional nervous system unit in its digestive system that acts in relative isolation from the rest of the nervous system, the pharyngeal nervous system. This simple nervous system serves as a prime model to investigate how neurons assemble into a functional circuit to produce behavior.

The pharynx, the core component of the foregut of all animals, is a myogenic organ that is innervated by an autonomously acting nervous system. In *C. elegans,* the pharynx contains 20 contractile myoepithelial-like cells, which pump bacteria from the environment into the intestine^6^. The pharyngeal nervous system of *C. elegans* consists of 20 neurons that fall into 14 anatomically distinct classes. 6 of these classes are each constituted by a bilaterally symmetric neuron pair (12 neurons total) while another 8 distinct neuron classes are defined by single, unpaired neurons (**Figure 1A**)^6^. In addition to the 20 muscle cells and 20 neurons, there are 9 epithelial, 9 marginal, and 5 gland cells. The pharynx can be described as a myoepithelial organ, as its contractile tissues exhibit both mesodermal and ectodermal properties^7^. Ensheathed by its own basal lamina and cuticle, the pharynx is separated from the remainder of the nervous system. Whereas in the somatic (non-pharyngeal) nervous system, the ectodermal nervous system is separated in the usual way from the mesodermal muscles by a basal lamina, in the pharynx no basal lamina separates motorneurons and the muscles they innervate. The pharyngeal and somatic nervous systems (defined as the entire nervous system with the exception of the pharyngeal nervous system^6^) are largely isolated from one another and are synaptically connected through only a single somatic neuron pair, RIPL/R^3,6^. Pharyngeal neurons can also communicate with the rest of the nervous system via non-synaptic actions of neuromodulatory biogenic amines and neuropeptides^8,9^.

**Figure 1.**
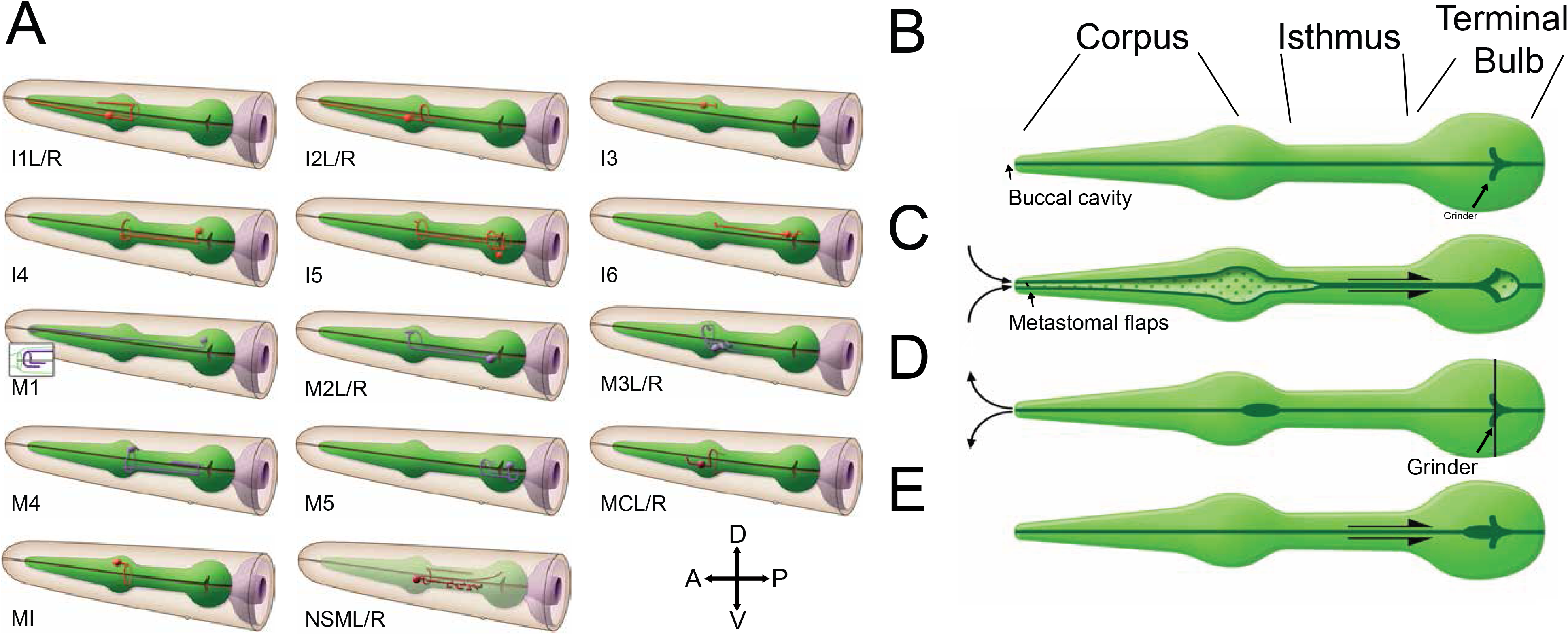
The *C. elegans* pharyngeal nervous system and feeding behavior. (**A**) Cartoon images of each pharyngeal neuron class used with permission from www.wormatlas.org. (**B**) The functional units of the pharynx: corpus, isthmus, and posterior bulb. Major sub-steps of feeding (top to bottom):(**C**) ingestion by the corpus, (**D**) fluid expulsion, and (**E**) isthmus peristalsis to deliver food to the grinder. After these steps, bacteria are passed through the pharyngeal-intestinal valve into the intestine.

The pharyngeal nervous system coordinates myogenic contractions in order to achieve filter feeding and regulates organ activity in response to environmental stimuli (Figure 1B). Because the worm’s body is held at a higher internal pressure than the environment, the pharynx functions as both a pump and valve. To feed, the worm first contracts the radial fibers of the corpus, anterior isthmus, and terminal bulb muscles to expand the lumen and pull in liquid and bacteria from the environment^10^. Bacteria are initially filtered based on size when moving past the buccal cavity and metastomal flaps^11^ (Figure 1C). A muscular relaxation event follows almost immediately after contraction, expelling liquid but trapping bacteria. Bacterial trapping happens through two complex and successive contraction and relaxation cycles, concentrating bacteria in the central lumen of the pharynx and expelling liquid via apical channels(Figure 1D)^11^. Following every 3-5 pumps, accumulated bacteria are advanced through the isthmus to the terminal bulb via peristaltic isthmus muscle contractions(Figure 1E)^12^. The grinder then crushes the bacteria before transport past the pharyngeal-intestinal valve into the intestine. Under optimal conditions a worm feeds at a neurogenic pumping rate of 3 Hz, which is entrained by pharyngeal neuron cholinergic signaling^13^. Touch^14^, sleep^15^, starvation^16^, light^17^, and odorant^18^ stimuli are capable of modulating the pumping rate. Despite detailed descriptions of pharyngeal behavior, the role the pharyngeal nervous system plays and the circuit details of how information is transmitted through the pharyngeal nervous system to generate behavior remains to be fully understood.

An anatomical atlas, including a synaptic wiring diagram of the *C. elegans* pharynx was first published in 1976^6^. However, while maps of individual neurons were generated, only cell-class connectivity data were published. Criteria for defining synapses have also evolved since these early reconstructions and so has the ability to analyze network structure computationally. Using modern day reconstruction methods we have recently reexamined the entire connectome of a *C. elegans* hermaphrodite and male. We reported the results of the reexamination of the somatic nervous system in the previous paper^5^ and we give here a detailed description of the pharyngeal connectome. Comparison of the pharyngeal nervous system to the somatic nervous system reveals its properties are quite distinct. Extending the previous analysis of the pharyngeal connectome^6^, we now describe connectivity of individual cells with anatomical weights. We delineate new synaptic in- and output for all pharyngeal neurons. We validate newly described connections using fluorescent reporter genes. Other novel observations include that most neurons possess sensory endings and that all neurons connect to end organs, allowing for local information processing in each functional domain of the pharynx, thereby revealing a shallow structure for information processing. We examine a number of distinct network features of the pharyngeal connectome and superimpose known molecular features onto the anatomical map. We speculate that the pharyngeal nervous system has features that may be characteristic of ancestral nervous systems.

## RESULTS

### Re-Analysis of the pharyngeal connectome

We have recently published a re-analysis of the synaptic connectivity of the entire *C. elegans* hermaphrodite nervous system as displayed in five legacy EM print series of five hermaphrodite animals (N2U, JSE, N2T, N2W, JSA)^5^. We describe here the salient features of synaptic connectivity within the pharyngeal nervous system, based on the N2T, N2W, and JSA series. The N2T and N2W series are largely non-overlapping and cover the anterior and posterior of the pharynx, respectively (Figure 2A). The JSA series is a biological replicate and overlaps the N2W series in the densely-connected region known as the pharyngeal nerve ring. The 20 neuronal, 20 muscle, 9 epithelial, 9 marginal, and 5 gland cells were reconstructed from over 1600 electron micrographs and were annotated for morphology and connectivity (**Supplemental data 1-2**). The most anterior (arcade cells and hyp1) and posterior (pharyngeal-intestinal valve) structures are not innervated by the pharyngeal nervous system and were not included in our analysis. We combined chemical synaptic and gap junction connectivity from N2T, JSA, and N2W (**Supplemental data 3**) and in the overlapping region of JSA and N2W we averaged the weight of connections. Connection weights were determined by summing the sizes of the usually multiple synapses involved in making the connection, which were determined by counting the number of EM sections traversed by each synaptic structure. Membranes were densely stained in the JSA series, and therefore we did not evaluate this sample for gap junction connectivity. We note that although the micrographs from which we reconstructed our datasets were generated using a chemical fixative rather than more modern high pressure freezing technology, preservation of certain ultrastructural anatomical features (e.g. presynaptic densities, neuronal gap junctions, sensory endings) remains excellent, as previously discussed^5,19^.

**Fig. 2.**
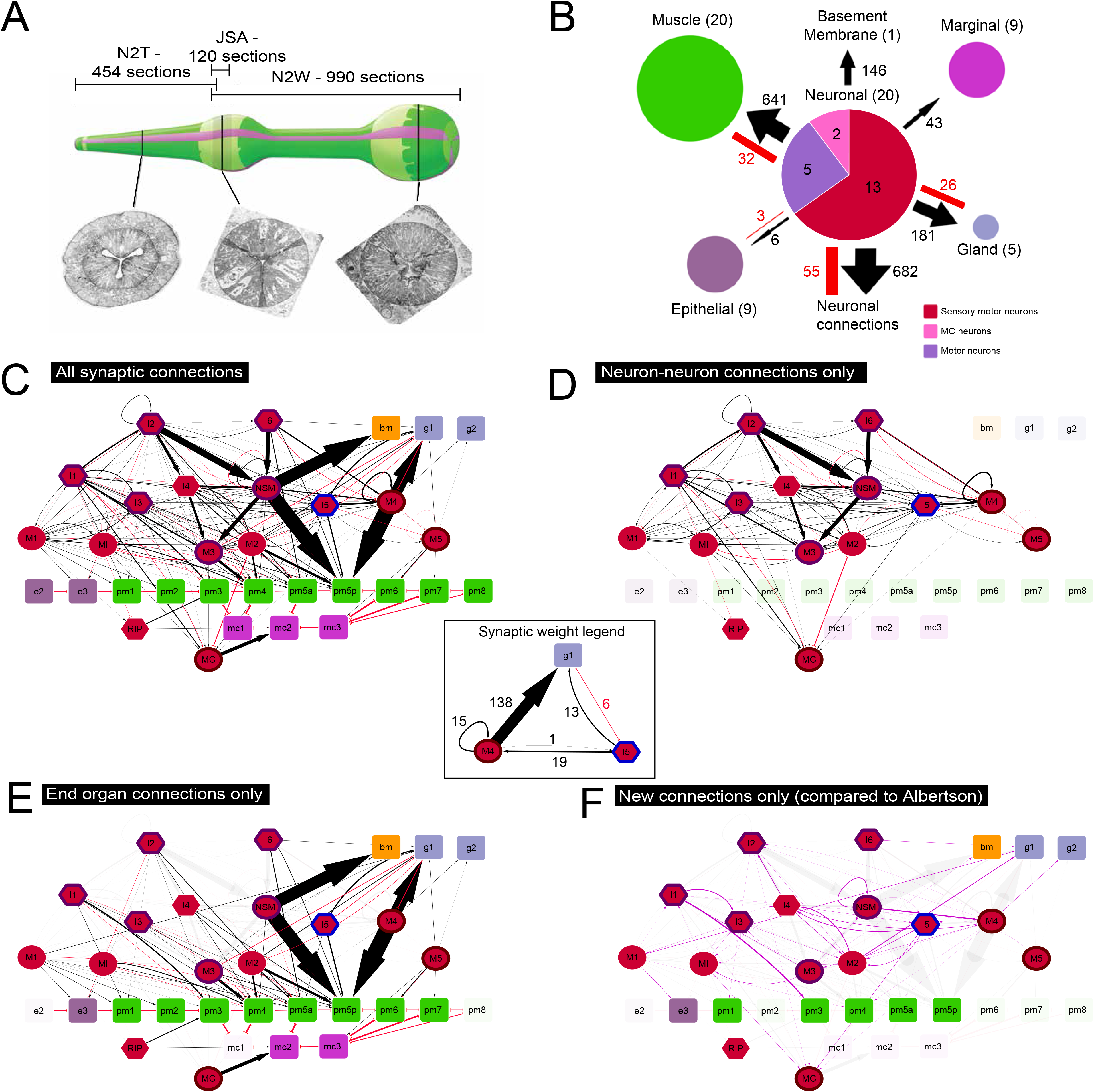
Circuits for feeding behavior. (**A**) Three electron micrograph series (N2T, JSA, N2W) were used to reconstruct the pharynx. The JSA and N2W series both cover the pharyngeal nerve ring (shown in gray hash pattern), which is the most complex pharyngeal neuropil. (**B**) Pharyngeal nervous system targets. Black arrows represent directed chemical edges and red lines represent undirected gap junction edges. Numbers in parenthesis are the number of individual cells per tissue. Numbers represent the synaptic weight (# serial sections). (**C**) Graphic layout of connected cell classes in the C. elegans pharynx. Square nodes are end-organs, including muscle (green), marginal (fuchsia), gland (blue bell), epithelial (deep pink), and basement membrane (orange). Interneurons are red hexagons and motorneurons are red circles. Neurons with outlines have either apical (purple), unexposed (brown), or embedded (blue) sensory endings. Directed chemical edges and undirected gap junction edges are represented by black arrows and red lines, respectively. The line width is proportional to the anatomical strength of that connection (# serial sections, see inset). All nodes represented are cell classes whose left/right or triradiate symmetry has been combined, except pm5 which was divided into its anterior (pm5a) and posterior (pm5p) components. (**D**) Diagram from C with only neuron-neuron connections highlighted. (**E**) Diagram from C with only connections to end-organs highlighted. (**F**) Diagram from C with only connections that were newly added by this reconstruction compared to Albertson et al 1976. Color codes for all panels match WormAtlas (www.wormatlas.org/colorcode.htm), except purple in (F) which indicates novel connections compared to Albertson et al 1976.

Our analysis resulted in a new and complete, weighted wiring diagram of the *C. elegans* pharynx (**Supplemental data 4, 5**)^5^. We generated a simplified layout of connectivity by cell class where anatomically equivalent cells are combined and shown as single nodes (Figure 2C). These cell classes were previously described by anatomical similarity, connectivity, and formaldehyde induced fluorescence staining^6^ and were consistent with the new connectivity data. To recapitulate the anatomy and the sequential flow of bacteria through the pharyngeal lumen, we arranged end-organ nodes according to their anteroposterior (A-P) position. Neurons were arranged vertically to reflect hierarchical patterns of synaptic connections. The pharyngeal connectome contains 1012 chemical and 96 electrical (gap junction) synapses. 495 of the chemical connections are neuromuscular junctions (NMJs)^5^. Of the chemical synapses, 486 (43.8%) are monadic, 585 (52.8%) are dyadic, and 38 (3.4%) are triadic. The somatic nervous system has a smaller proportion of monadic (35%), similar proportion of dyadic (54%), and larger proportion of triadic (10%) synapses. We also assessed morphology of synapses, allowing a more accurate prediction of anatomical connection strength. Our synaptic weight metric (#serial EM sections), a proxy for anatomical strength, was calculated by summing the number of serial sections with synaptic specializations. This synaptic weight metric has a large range, varying from 1 to 78 for individual connections between neurons. We generated maps with connectivity for each cell and identified many connections not previously noted^6^. Several neuron classes previously reported to be purely presynaptic do indeed receive synaptic input, yielding a network where all neurons make and receive chemical synapses as well as gap junctions. We were able to verify 94% (50/53) of the chemical synaptic connections between cell classes previously reported in the pharynx^6,20^. The three discrepant connections are small (two or fewer synapses), and we observed the reciprocal connection in two of the three instances. Our pharyngeal reconstruction contains 142 chemical connections between cell classes, 50 of which were not originally reported (Figure 2F). We believe that the primary reason for this discrepancy is technological advancement since our digital reconstruction and database curation allowed us to more accurately score and transcribe synapses. These new connections ranged in strength (1 to 25.5 sections of connectivity) with a mean and median sizes of 4.46 and 3 sections, respectively. Their total weight is 190 sections, representing 12.1% of the total chemical synaptic network. 38/95 novel connections are made onto end organs, including 19 NMJ connections.

The only direct synaptic connections with the somatic nervous system are through the pair of non-pharyngeal RIPL/R neurons. The RIP neurons have cell bodies in the somatic anterior ganglion, dendritic processes which enter the somatic nerve ring, and anterior axons with project toward the nose and cross the pharyngeal basal lamina and where they make gap junctions with the pharyngeal neuron I1. We also observed a previously unreported synapse from the M1 motorneuron onto RIP in the N2T series (Figure 2C), as well as synapses from the RIP axon onto anterior pharyngeal muscles.

### Fluorescent active zone reporters confirm novel synaptic patterns

Our new reconstruction revealed at least one previously unreported synaptic connection for each pharyngeal neuron class (Figure 2F). In addition to new connections between cells, we also found new locations of synaptic output within the pharyngeal nervous system (red arrows). To confirm these new synapses in live animals, we created transgenic lines expressing fluorescently-tagged synaptic proteins (Figure 3). We first confirmed that tagged RAB-3, a vesicle associated GTPase, and tagged CLA-1, the functional homolog of the active zone proteins Piccolo and fife^21^, were both subcellularly distributed similar to reconstructed electron micrographs (blue arrows). We expressed tagged RAB-3 (Figure 3A,C) and tagged CLA-1 (Figure 3B,D) in the NSM and I1 neurons and observed fluorescent puncta at presynaptic locations predicted by EM, with GFP::CLA-1 showing a more punctate pattern(Figure 3B,3D). We compared our results for NSM->M3 synapses and found a good correspondence between fluorescent synaptic puncta and EM synapses (Figure 3E-F). We then quantified presynaptic active zones by counting fluorescent CLA-1 puncta along the length of the I2, I1, MC, and I5 neurons in independent transgenic lines. Nearly half of pharyngeal neuron classes (I1, I2, I3, I6, M3, MC) were originally described to exhibit an axodendritic separation between their two neurites. Our reconstruction, as well as an independent reconstruction of the I2 neuron^17^, revealed that several bipolar pharyngeal neurons make synaptic output on both of their neurites. We confirmed the presence of active zones on both neurites in the I2 (Figure 3G-I) and I1 (Figure 3J-K) neurons.

**Fig 3.**
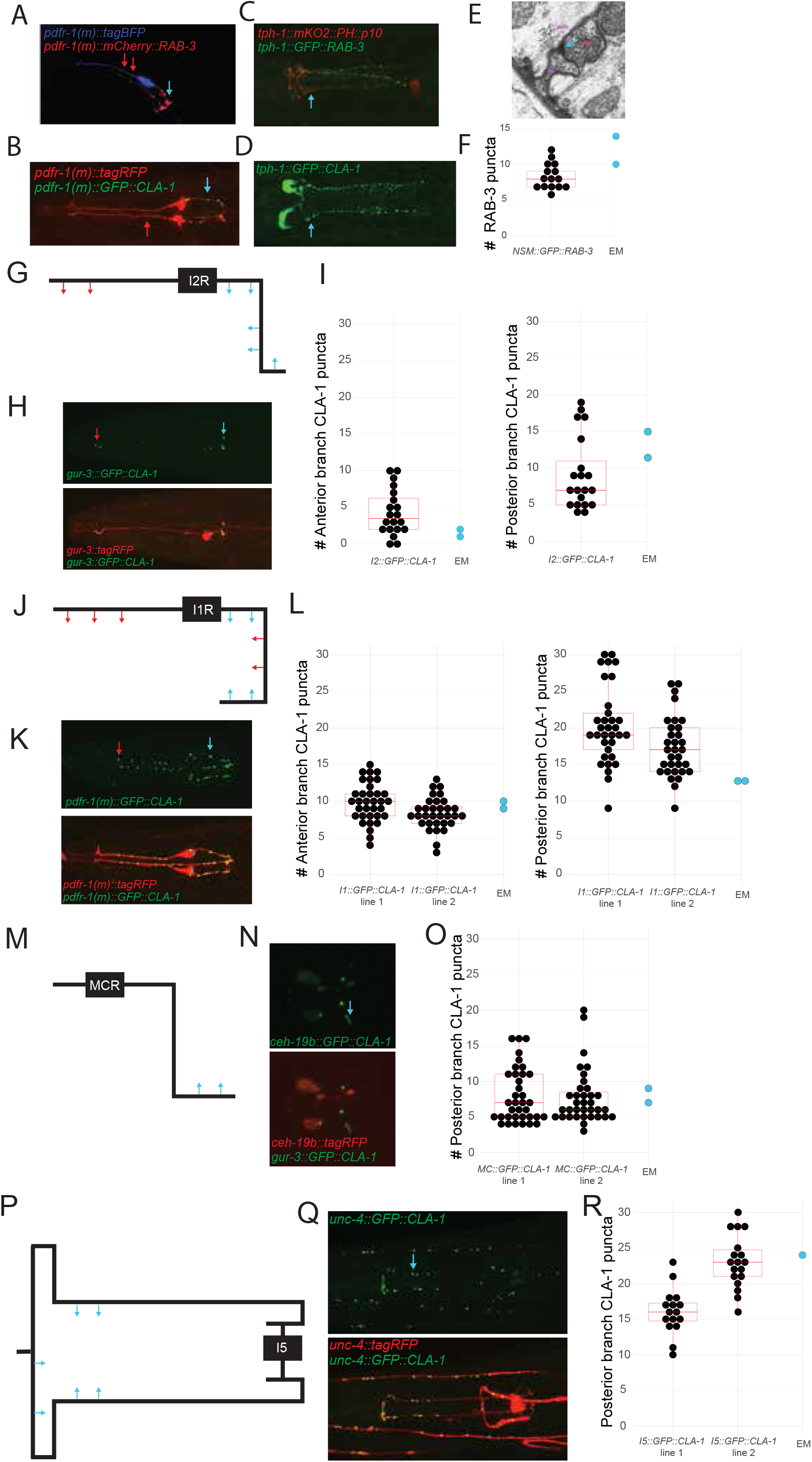
Confirmation of ultrastructural connectivity with fluorescent active zone reporters. (**A**) Maximum intensity projection of I1 synapses and cytoplasm labeled by RAB-3 and mTagBFP2. (**B**) Maximum intensity projection of I1 synapses and cytoplasm labeled by CLA-1 and tagRFP. (**C**) Maximum intensity projection of NSM synapses and M3 cytoplasm labeled by RAB-3 and mKO2. (**D**) Maximum intensity projection of NSM synapses labeled by CLA-1. (**E**) Example EM image showing NSM synapse (red star) adjacent to M3.(**F**). Comparison is made to EM observations with quantification of C, counting RAB-3 puncta adjacent to the M3 cell.. (**G**) Schematic of reconstructed I2R neuron, showing anterior branch (left of black cell body), posterior branch (right of black cell body), and locations of active zones (blue designates previously reported active zones (Albertson and Thomson 1976), red designates those reported in this study). (**H**) Maximum intensity projection fluorescent images of *gur-3::GFP::CLA-1* and *gur-3::tagRFP.* (**I**) Quantification of CLA-1 puncta in the anterior (left) and posterior (right) branches compared to EM observations. (**J**) Schematic of the I1R neuron. (**K**) Maximum intensity projection fluorescent images of *pdfr-1(m)::GFP::CLA-1* and *pdfr-1(m)::tagRFP*. (**L**) Quantification of CLA-1 puncta in the anterior (left) and posterior (right) branches compared to EM observations. (**M**) Schematic of the rich MC neuron. (**N**) Maximum intensity projection fluorescent images of *ceh-19(b)::GFP::CLA-1* and *ceh-19(b)::tagRFP*. (**O**) Quantification of CLA-1 puncta in the posterior branch compared to EM observations. (**P**) Schematic of the I5 neuron. (**Q**) Maximum intensity projection fluorescent images of *unc-4::GFP::CLA-1* and *unc-4::tagRFP*. (**R**) Quantification of CLA-1 puncta compared to EM observations.

Neurons such as MC and I5 have spatially restricted synaptic output, suggesting compartmentalization of synaptic zones along the length of a neurite. We confirmed spatially restricted active zones along the posterior axon of MC (Figure 3M-O) and anterior axon of I5 (Figure 3P-R). Moreover, in all reporter lines evaluated, we found that the number of synapses we observed in our reconstructions (blue dots) were within the range of observed synaptic reporters. Together these results suggest that pharyngeal neurons exhibit more complex synaptic patterns than previously realized and reject assumptions of axonal and dendritic branches. Despite variability in the number of synapses observed per animal, there was a strong conservation in the subcellular distribution of synaptic puncta.

### All pharyngeal neurons target end organs

Our analysis of the pharyngeal wiring diagram revealed a number of intriguing and previously underappreciated features. All non-neuronal tissues within the pharynx receive synaptic input and can be classified as end-organs (Figure 2B). The total chemical synaptic output directed toward end-organs (59.9%) is greater than connectivity between neurons (40.1%) (Figure 2B,D, E). The most frequent non-neuronal synaptic target is muscle, with 13/14 pharyngeal neuron classes making muscle output. The outlier is the MC neuron, which has been electrophysiologically shown to function as a motorneuron^22^, yet synapses solely onto marginal cells. Eight pharyngeal neurons synapse onto gland cells, making it the second most-heavily innervated end organ. The M4 motorneuron, which makes frequent dyadic synapses onto muscle and gland, makes more NMJs than any other pharyngeal neuron. M4’s output onto gland and muscle cells suggests a potential relationship between peristalsis and gastric enzyme release, perhaps coupling digestion to pharyngeal activity. Another large pharyngeal connection is NSM’s serotonergic synapse across the pharyngeal basal lamina into the pseudocoelom. The NSM neurons are a major source of serotonin to the whole animal and have been shown to modulate the slowing response of the worm upon encountering a food patch^8^. Epithelial cells inside the pharynx are also innervated. These cells contain short radial intermediate filaments anchored to both the apical and basal membranes, providing structural integrity to the pharynx during vigorous movement of the whole organ, but are not known to be contractile. Taken together, the network architecture of the pharyngeal nervous system appears remarkably shallow with all neurons directly targeting effector tissue.

### Sensory endings are distributed throughout the pharynx

In our reconstruction we found not only new synaptic connections and many previously unreported end organ connections, but also discovered apparent sensory endings of many pharyngeal neurons that are distributed along the entire pharynx (Figure 4A). Examination of cellular morphology revealed 15 out of the 20 pharyngeal neurons (10 out of 14 classes) with putative sensory function (Figure 2B, Figure 4B). Three types of ultrastructural specializations are present, classified as: apical (Figure 4C), unexposed (Figure 4D), and embedded (Figure 4E). Pharyngeal sensory endings are primitive in appearance, and do not include basal bodies, Y-links, axonemes, or cilia^23–25^. Sensory structures consist of small neuronal projections that are anchored by adherens junctions to the cuticle or lumen at critical locations where bacteria pass through the pharyngeal lumen. Pumping module neurons I1, I2, and I3 form apical sensors with the pharyngeal lumen to detect bacteria near the metastomal flaps. Other neurons with apical sensors include NSM near the sieve (**Supplemental Figures 1 and 2**), M3 at the anterior isthmus, and I6 in the terminal bulb. The unexposed sensors of the pharynx are putative internal proprioceptive endings that may measure muscle movement. MC and M4 form unexposed sensors near the sieve and isthmus, respectively. M5 forms an unexposed putative sensor close to the terminal bulb’s lumen. I5 has the only putative sensor embedded within a pharyngeal muscle located near the grinder. Visualized along the A-P axis of the pharynx, putative sensors are localized to detect sequential bacteria travel through the pharyngeal lumen during feeding (Figure 4A).

**Fig. 4.**
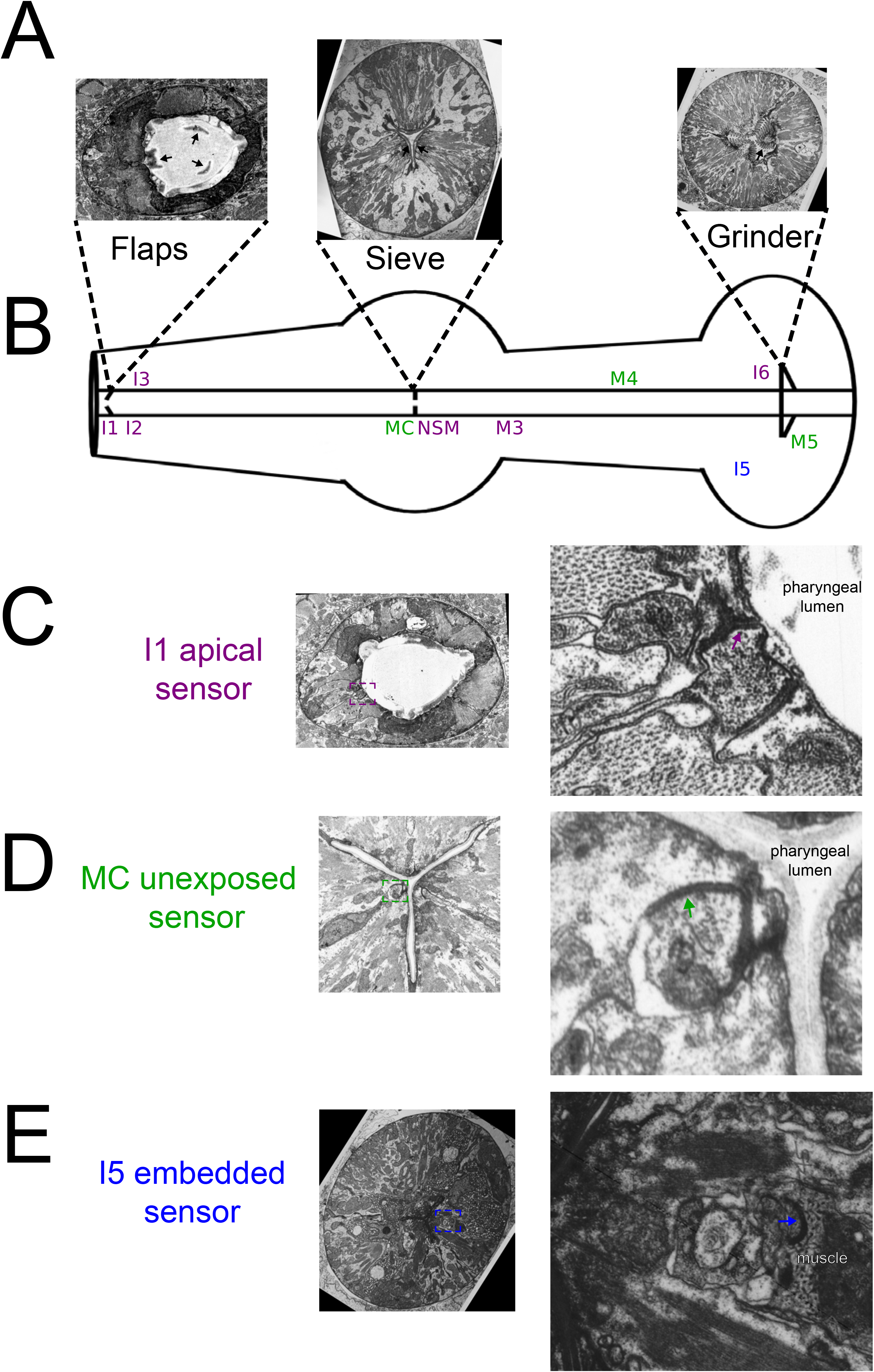
Multiple types of putative mechanosensory endings are distributed throughout the pharynx. (**A**) Structures for physically separating and processing bacteria include the flaps, sieve, and grinder. (**B**) Positions of apical (purple), unexposed (green), and embedded (blue) sensors are shown symbolically by neuron name along the length of the pharynx, relative to their A/P locales along the pharyngeal lumen (except I5 with an embedded sensor). All lie close to the pharyngeal lumen except for I5. (**C**) Example of an apical (exposed) sensor of the I1 neuron shown near the flaps. (**D**) Example of an unexposed sensor of the MC neuron shown near the sieve. (**E**) Example of an embedded sensor of the I5 neuron shown near the grinder. Short arrows in right panels indicate adherens junctions.

Exposed sensory endings make direct contact to the internal pharyngeal cuticle, very close to a cuticular specialization, so that luminal contents (bacteria) might be directly sensed, or deflections of the cuticle specialization (flap, sieve, grinder tooth, or isthmus) will be detected. Unexposed endings also lie close to the cuticle, but are shielded by a thin wrapping of cytoplasm from neighboring pharyngeal myoepithelial cells. They may sense the presence of absence of bacterial contents in the lumen through larger deflections of cuticle specializations, or by proprioception of muscle contraction. Embedded sensory endings lie much deeper in the pharyngeal tissue, far from the lumen, and are likely to respond to local pressure or stretching of the muscle tissue (e.g. proprioception) during radial contractions. The cytoplasm within these short branches often contains clusters of small vesicles and short cytoskeletal fibrils, but no organized axoneme^23^. The sensory branchlet is often irregular in shape but stout, firmly connecting to surrounding cells by thick adherens junctions around the entire circumference of the branch. Taken together with our finding that most neurons directly connect to end organs, our analysis should lead to a re-evaluation of how *C. elegans* pharyngeal neurons are classified. Because of the oversight of many NMJs, the pharynx was originally classified to contain 6 classes of interneurons, whose names begin with an ‘I’, and 7 classes of motorneurons beginning with an ‘M’6. Our connectivity data suggest the pharyngeal nervous system has no true interneurons but is instead mostly comprised of multi-functional sensory-motorneurons. The predominance of sensory-motorneurons underscores the unexpectedly shallow information processing depth within the pharyngeal nervous system.

### The pharyngeal connectome is organized in functional domains and lacks apparent network motifs found in the somatic nervous system

We next asked if the shallow pharyngeal connectome contains groups of cells dedicated to a specific sub-behavior and evaluated the network for modular structural components. We applied the Louvain Method for community detection^26^ to the combined the weighted chemical synaptic + gap junction networks to identify groups of cells more strongly connected to each other than other cells in the network. For this analysis, we assumed that the functional strengths of chemical and gap junction connections, based on the morphological sizes, were equivalent. This method generated a partitioning with four functionally relevant modules (Q = 0.352). We named these modules for their primary inferred behaviors supported by functions of individual neurons within module: pumping, neuromodulation/relaxation, peristalsis, and grinding (Figure 5A-D). The individual neurons in each module either have a consistent functional description or have no experimentally defined function (**Supplemental data 6**). For those neurons with unknown function, we can predict an association with the following behaviors based on module identity: pumping (I3, MI), neuromodulation/relaxation (I4, I6), and grinding (M5). While the grinding module assignment is the most tenuous, it has been shown that the pharyngeal nervous system, presumably in part through M5, controls terminal bulb contraction^13^. It should be noted that our method for community detection produces discrete partitioning, and thus the M3 neurons segregated into different modules due to differences in their connectivity across the left/right axis (**Supplemental data 6**). Three modules also directly overlay in anatomical position with the three functional domains of the pharynx (pumping module with corpus, peristalsis module with isthmus, and grinding module with terminal bulb). The neuromodulation/relaxation module neurons target the entire pharynx. In conclusion, as all modules contain both sensory specializations and motor output, each functional domain of the pharynx can sense its local environment and modulate organ function as bacterial and muscle activity travel along the A-P axis of the pharynx.

**Fig. 5.**
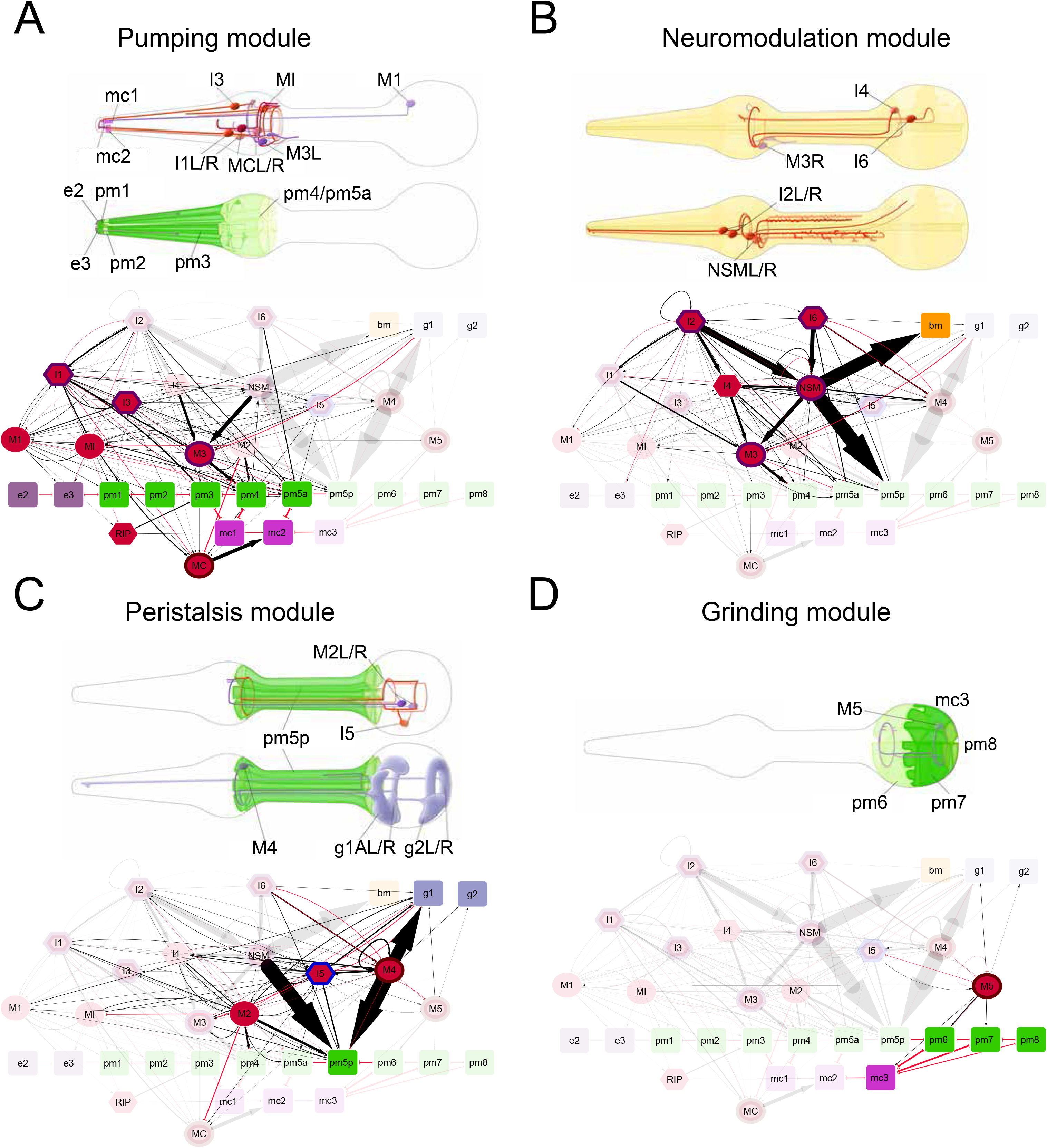
Computational modules overlay with functional units for feeding behavior. (**A**) The anterior pumping module. The pumping-rate controlling MC neurons, and the marginal cells they innervate, are present. This module contains the only somatic nervous system connections, connecting to the RIP neurons, which are also necessary for controlling pumping-rate off food. For clarity, M3 neuron class is shown in this module. (**B**) The neuromodulation/relaxation module. The serotonergic neurosecretory neuron NSM and glutamatergic relaxation promoting M3R neuron are members. The I2 neuron has also been shown to directly sense the environment and inhibit pumping in a monosynaptic circuit^17^. NSM downstream targets include many members of the somatic nervous system (not shown). (**C**) Peristalsis module. The M4 neuron, essential for peristalsis, also makes the largest NMJ in the pharynx (M4->pm5). All gland cells of the pharynx are also present, suggesting a potential role in digestive activity and/or molting. (**D**) Grinding module. M5, the single neuron in this module, is the only cell in C. elegans to innervate the pm6 and pm7 muscles. Colors for all tissues can be found at https://www.wormatlas.org/colorcode.htm.

Another way anatomical connectivity contributes to network properties is through over-represented patterns, or motifs. In a variety of biological systems, motifs have been implicated as building blocks of networks^27^. We observed two-neuron and three-neuron motifs of chemical connectivity and compared their frequency to randomized networks with similar structure. We found that the pharyngeal chemical synaptic network contains no statistically significant over-represented two-neuron motifs (Figure 6A), while the somatic nervous system had two of three motifs statistically over-represented (Figure 6B). For three-neuron motifs, we found that only two are statistically over-represented in the pharyngeal nervous system (Figure 6C) compared to eight somatically (Figure 6D). Furthermore, some three-neuron motifs found in both the somatic nervous system and our randomizations were not observed in the pharyngeal nervous system (Figure 6C-D). These results corroborate the shallowness of the pharyngeal network and reinforce how its overall structure is distinct from the *C. elegans* somatic nervous system.

**Fig 6.**
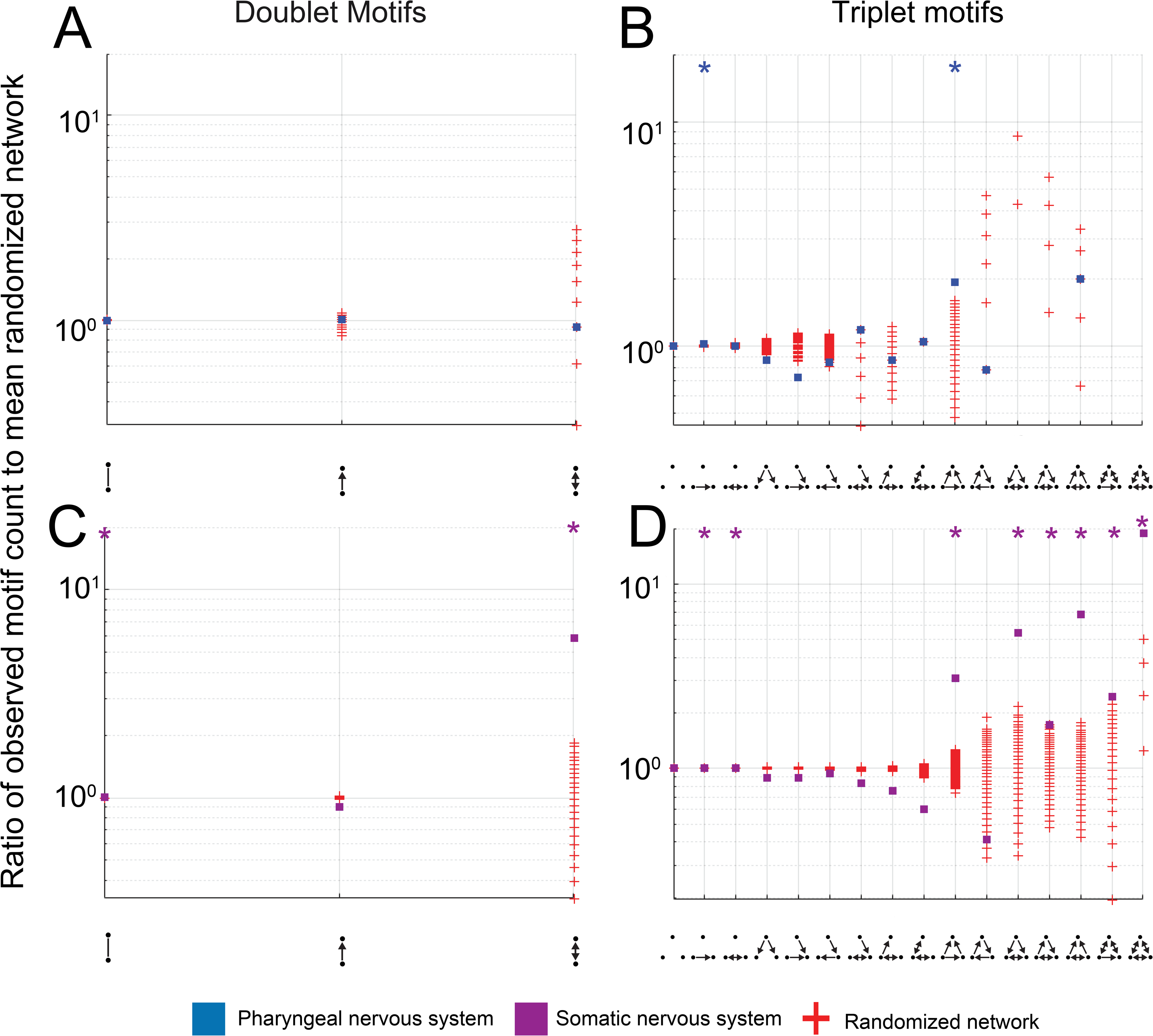
Network motif analysis. Occurrence of doublet and triplet motifs of the pharyngeal and somatic chemical synaptic networks. (A) Doublet and (B) Triplet motifs for the pharyngeal nervous system. (C) Doublet and (D) triplet motifs for the somatic nervous system. Plotted squares dots represent the ratio of observed doublet and triplet motifs to an average obtained from 1000 randomized networks with preserved network properties. Randomized networks are plotted as a red ‘+’. Absence of a square occurs when that network motif was not observed within the connectivity data. Motifs that are statistically overrepresented compared to randomized data were calculated using the single step min p procedure and multiple hypothesis testing (* = p =< 0.0005).

### Small connections predominate in the pharyngeal connectome

Pharyngeal neurons connect to multiple cell types over a range of synaptic strengths. We compared the distributions of in degree (number of incoming edges) and out degree (number of outgoing edges) for the chemical synaptic networks of the pharyngeal (blue) and somatic (purple) nervous systems and found them both to be significantly different (KS=0.438, p=2.207×10^−8^ and KS=0.358, p=9.22×10^−6^, respectively) (Figure 7A).The distribution of neighbor number (gap junction edges are undirected) was also different between the pharyngeal and somatic nervous systems (KS=0.224, p=0.011) (Figure 7B). Beyond number of connections, we compared the size of connections for the chemical and gap junction networks and found that both have different distributions (KS=0.110, p=0.005 and KS=0.409, p=8.01×10^−18^, respectively) (Figure 7C-D). Small edges may be important to network structure, as they comprise a considerable amount of the total load through the network, so we compared the cumulative load (sum of all synaptic weights) distribution for both networks and found a statistically significant difference for chemical synapses (KS=0.484, p=3.09×10^−5^) but not gap junctions (KS=0.487, p=0.076) (Figure 7E-F). In conclusion, the pharyngeal neurons connect to fewer partners with small synapses, on average.

**Fig. 7.**
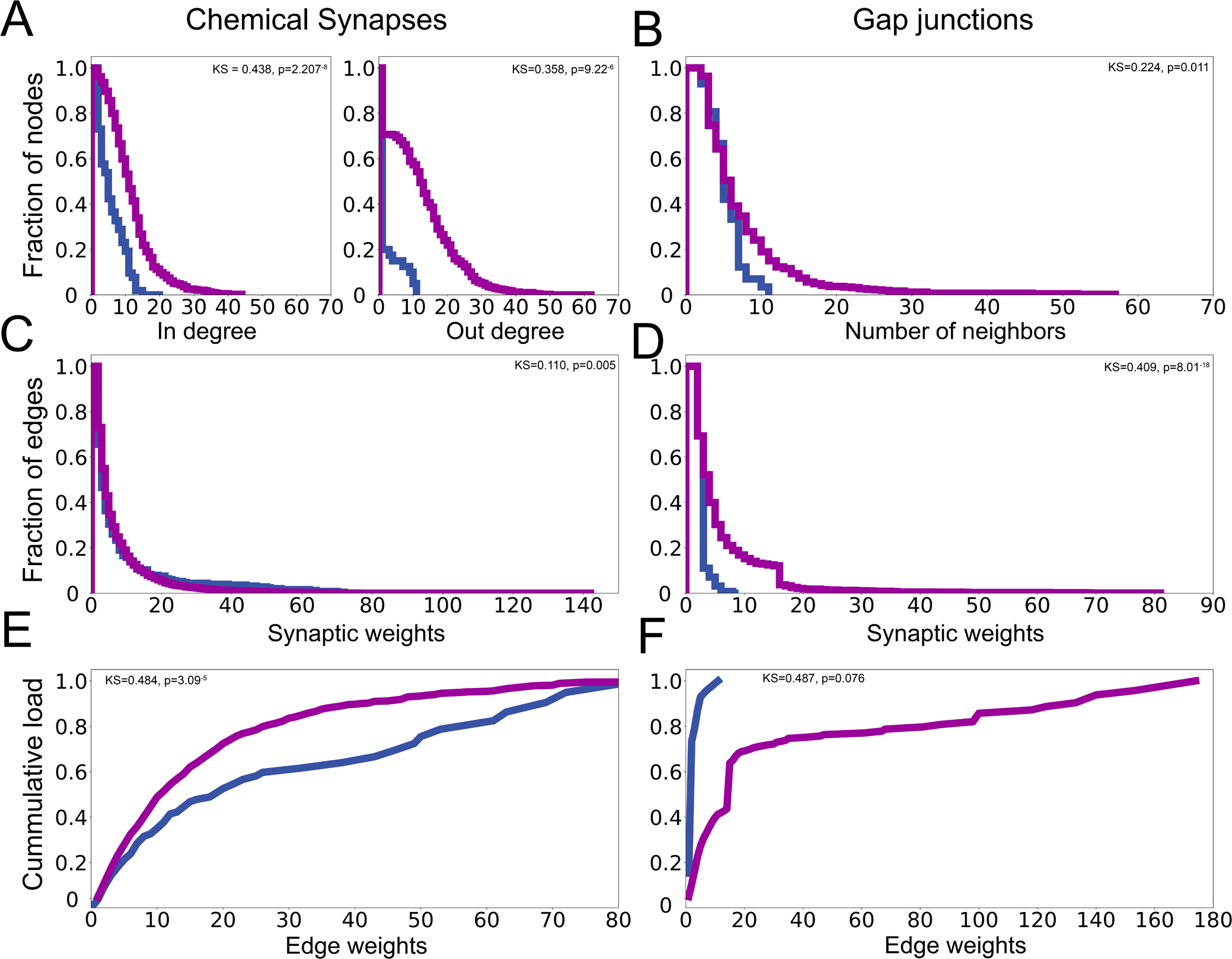
Pharyngeal and somatic connectivity networks have different structural properties. (**A**) In Degree distributions and out degree distributions for chemical synaptic networks of the pharyngeal (blue) and somatic (green) nervous systems. (**B**) Degree distributions for gap junction connectivity. (**C**) Distribution of synaptic weights for chemical synaptic and (**D**) gap junction networks. Synaptic weight is calculated by summing the number of individual 70- to 90-nm serial sections where a presynaptic specialization is observed. (**E**) Cumulative load distribution through chemical synaptic and (**F**) gap junction networks, calculated by summing all edge weights. Distributions in A-F were compared by using a two-sample Kolmogorov-Smirnov test with two-tailed p-value.

### Pharyngeal synapses are variable within and between animals

Small synaptic connections can vary within and between individuals^28,29^. To determine the amount of variability within a single individual, we compared wiring differences between presumptively equivalent homologous left/right neuron pairs. We analyzed the chemical synaptic output of six neuron classes in each sample that have left/right homologous pairs (I1, I2, M2, M3, and MC) and found that only 35 of 83 chemical edges (42.1%) are present on both sides of the animal, revealing a large amount of intraindividual variability. To determine whether network flow is biased toward one side of the animal we evaluated the average size and total load of synaptic connections through each side of the network. There was also no statistical difference in average size of synaptic output across the left/right axis (6 vs. 4.18 sections, unpaired t-test, t=1.61, df=81, p=0.111288). Of note is the inhibitory glutamatergic motorneuron M3, where in both reconstructions the left M3 neuron (M3L) makes over twice as many synapses as the right M3 neuron (M3R), including synapses to distinct postsynaptic partners. The different connectivity patterns of M3L and M3R place each neuron into distinct functional modules (**Fig.5**). This result contrasts with the somatic nervous system, where left/right homologous neurons typically make similar connections^4,5^.

The availability of two reconstructions (N2W and JSA series) covering the pharyngeal nerve ring allowed us to evaluate inter-individual differences in connectivity. In the somatic nervous system, detailed inter-animals comparisons of synaptic connectivity have only been completed between animals of different developmental stage or sex. We compared chemical synaptic connectivity of the pharyngeal nerve ring between age and sex matched controls, allowing us to evaluate the reproducibility of connectivity for all neurons except I6 and M5. There are 30 connected cells common to each reconstruction. These cells are connected by 145 chemical edges in JSA, 108 chemical edges in N2W and 59 chemical edges present in both series, with 86 and 49 connections specific to JSA and N2W, respectively (Figure 8A). A cumulative density function for both common and unique edges shows that at all synaptic weights, the edges unique to the JSA series are smaller, on average (Figure 8B). Common edges have a mean size of 6.74 sections, significantly different from the 4.85 section average size for edges unique to only one sample (unpaired t-test, t=2.0102, df=192, p = 0.045809). This trend is common among *C. elegans* EM comparisons. On average, synaptic connections common across developmental stages or sex are larger than unique connections^29,30^. Despite being sex- and age-matched biological replicates, the pharyngeal connectivity of the N2W and JSA series still showed many differences in individual connections. Together, these intra- and interindividual comparisons indicate the pharyngeal wiring diagram is more variable than previously reported^28^.

**Fig. 8.**
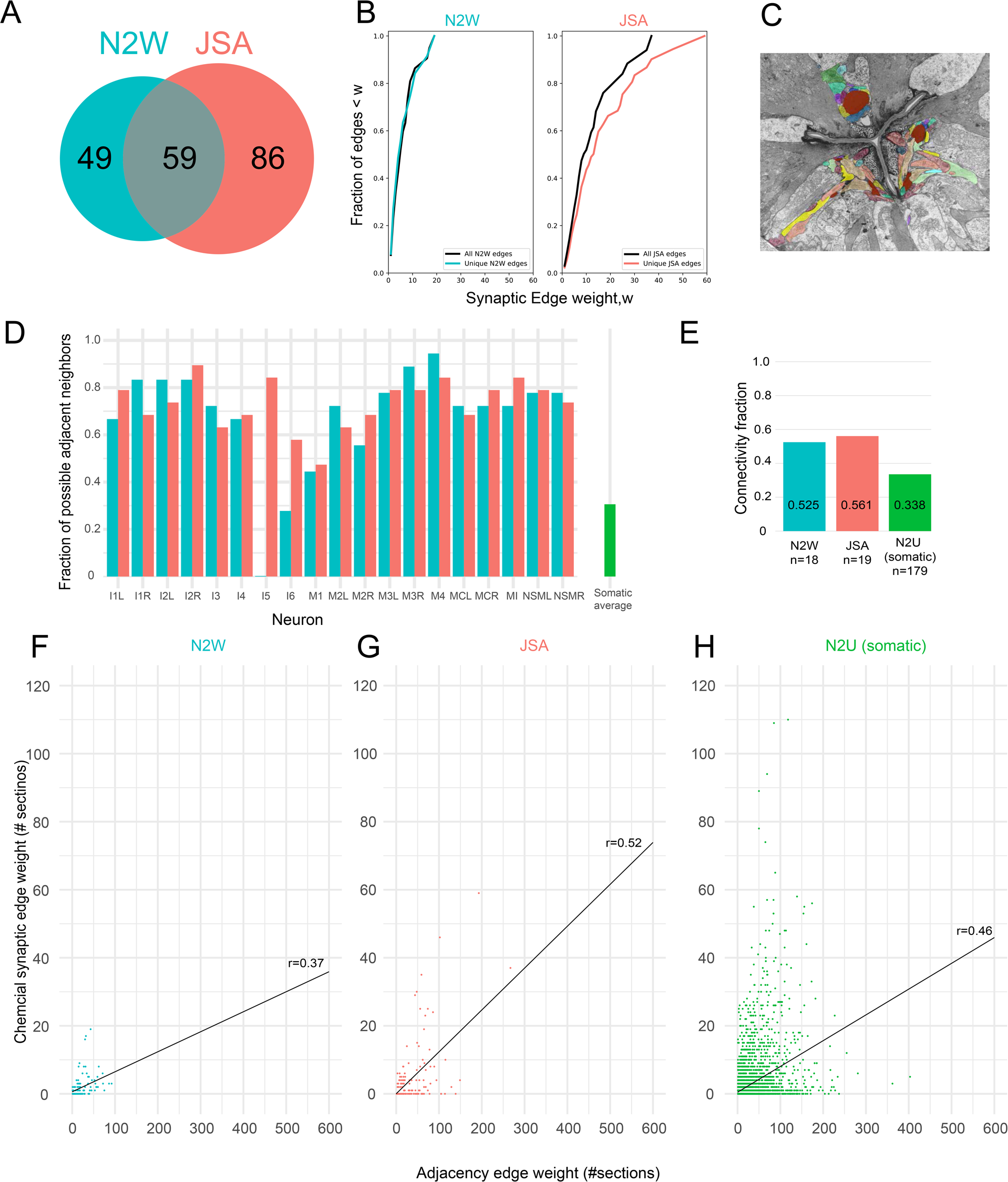
Comparison of unique and shared edges between replicate nerve ring reconstructions. (**A**) Overlap in chemical connections between N2W and JSA reconstructions. (**B**) Cumulative density function of two replicate pharyngeal nerve ring reconstructions, N2W (left) and JSA (right). The distributions of edges common to both series are in black, and those unique to N2W and JSA in teal and red, respectively. (**C**) Example image of volumetric reconstruction of neuron profiles in the JSA nerve ring. (**D**) Fraction of possible adjacent neighbors plotted for each neuron in the N2W (teal bars) and JSA (red bars) with the average of the adult somatic nerve ring in green. (**E**) Connectivity fraction (undirected chemical edges divided by undirected adjacency edges) for the N2W, JSA, and N2U (adult somatic nerve ring) series, n=number of neurons within series. (**F**) Adjacency edge weight vs chemical edge weight plotted for the N2W series with regression line plotted in black. (**G**) Adjacency edge weight vs chemical edge weight plotted for the JSA series with regression line plotted in black. (**H**) Adjacency edge weight vs chemical edge weight plotted for the N2U (somatic) series with regression line plotted in black. Spearman’s correlation coefficients are shown for each series.

### Correlation of synaptic targeting choice with neurite neighborhood

To form proper *en passant* synaptic connections, the processes of neurons must first be placed in a specific relative neighborhood to physically contact their synaptic partners. To evaluate the stereotypy of process placement as well as the relationship between neuronal process neighborhood and synaptic connectivity, we volumetrically reconstructed the pharyngeal nerve ring of both the N2W and JSA series (Figure 8C). We computationally extracted all neuronal neighbors from our reconstructions as previously described^30^, revealing 7820 individual neuronal adjacencies (adjacency defined as two touching membranes within any location of an EM section) (**Supplemental data 7**). Of the theoretically possible 171 adjacencies (19 choose 2), we observed 132 and 116 adjacency edges in JSA and N2W series, respectively (**Supplemental data 8**). The M5 neuron is the only pharyngeal neuron absent from the pharyngeal nerve ring and is therefore not included in our volumetric analyses. Within the pharyngeal nerve ring, each neuron is on average adjacent to 14 other neurons (range of 0 to 18)(Figure 8D). This translates to an average of 72.3% of all possible neuron neighboring each other, more than double the average of 30.2% in the adult somatic nerve ring. With the exceptions of I5 and I6, we saw good agreement in the number of adjacent neighbors in each series. The I5 neuron is absent from the N2W pharyngeal nerve ring, and the I6 neuron projects farther anterior in the JSA series, leading to the largest inter-animal discrepancies. Compared to chemical synaptic connectivity, adjacency edges are more conserved, with 102 edges present in both series, 30 JSA-specific (21 of which include I5 and/or I6), and 14 N2W-specific.

We also found that the connectivity fraction (number of synapses / number of adjacent neighbors) for the pharynx is larger than in the somatic nervous system. The connectivity fractions for the N2W and JSA series are 52.5% (61 / 116) and 56.1 (74 / 132), respectively. This connectivity fraction is greater than the somatic nerve ring 33.8% (1812 / 5368), demonstrating that not only do pharyngeal neurons contact more neighbors they also create synapses with them more frequently (Figure 8E). To further probe this issue, we asked whether synapses which are discrepant between our reconstructions were due to missing neuronal adjacencies. Of the 49 synaptic connections observed in N2W but not JSA, 49% (24/49) are neighboring neurons. 41.3% (36/87) of synaptic connections observed in JSA but not N2W are neighboring neurons, 24.1% (36/87) are impossible due to the lack of I5 in the N2W pharyngeal nerve ring. These discrepancies are likely not due to a proclivity of pharyngeal neurons to synapse randomly onto adjacent neighbors, as 24.5% (25/102) of identical adjacency edges across species are not synaptically connected.

We lastly asked whether the weight of contact is correlated with the weight of connectivity (#serial sections). Both weight metrics we measured for adjacency (# adjacent sections and #pixels where two processes are adjacent) are highly correlated (Spearman’s rho=0.93, p < 2.2 × 10^−16^ and 0.93, p < 2.2 × 10^−16^) in both EM series. The correlation between adjacency (#EM sections) and chemical connectivity (#EM sections) is low in both series (N2W spearman’s rho = 0.52, p= 1.15 × 10^−9^)(Figure 8F) (JSA spearman’s rho = 0.37, p= 1.37 × 10^−5^) (Figure 8G). A similar correlation was found for the adult somatic nerve ring (N2U spearman’s rho = 0.46, p=0.0000) (Figure 8H). Our volumetric analyses therefore shows that pharyngeal neurons not only physically contact more of their possible partners but also create synapses with a greater proportion of their neighbors. Nevertheless, there are outliers to this trend, such as the MC neuron which contacts between 13 and 15 neurons, yet synapses solely onto the marginal cell. Hence, the extent of contact between two neurons is not an entirely sufficient predictor for connectivity or strength of connectivity.

### Network analysis as a tool to predict neuronal function

How important are individual neurons to pharyngeal function? Traditionally, cell ablation experiments have been used to determine the importance of individual or sets of neurons to pharyngeal function. With the exception of M4, the neuron essential for isthmus peristalsis, all neurons can be killed without impairing the worm’s ability to survive under laboratory conditions. In the absence of M4, the posterior isthmus muscles cannot contract and the worm does not feed. When ablated, some neurons yielded a strong phenotype where pharynx that was deficient in pumping^31^. Two important caveats of this previous analysis are that 1) neurons were removed in early larval stages, and post-operative compensatory rewiring processes, previously observed to occur in the worm^32^, cannot be excluded and 2) sample size for embryonic ablations was very small. It remains clear that the pharyngeal nervous system is robust toward perturbations, and/or may have compensatory mechanisms for cell loss. In view of this robustness, we asked which neurons may be important for the function of the entire network by analyzing betweenness and closeness centralities. Betweenness centrality is a measure of how frequently a node is used in the shortest path between other nodes, while closeness centrality measures how long it will take information to diffuse from one node to all other nodes in the network. Overall, we observed a moderately strong relationship between the betweenness centrality and closeness centrality values, especially for left/right homologous neuron pairs (Figure 9). We also found a group of neurons that have high scores for both betweenness and closeness centrality. Consistent with the strongest ablation phenotype in the pharynx^31^, our centrality analysis identified M4 as one of the most centrally important pharyngeal neurons. Two other central neurons, I2 and M1, have recently been shown to promote light induced pumping inhibition^17^ and spitting behaviors^33^, respectively. We found previously unreported NMJs made by both of these neurons, which supports experimental findings that they act as functional monosynaptic circuits^17,33^. Interestingly, the I5 and M5 neurons have high betweenness and closeness centrality scores, implicating an important network function despite lacking strong ablation phenotypes such as abrogation of pumping^31^. Our centrality approach suggests functionally important neurons, as well as predict network importance for neurons whose function is not fully understood.

**Fig. 9.**
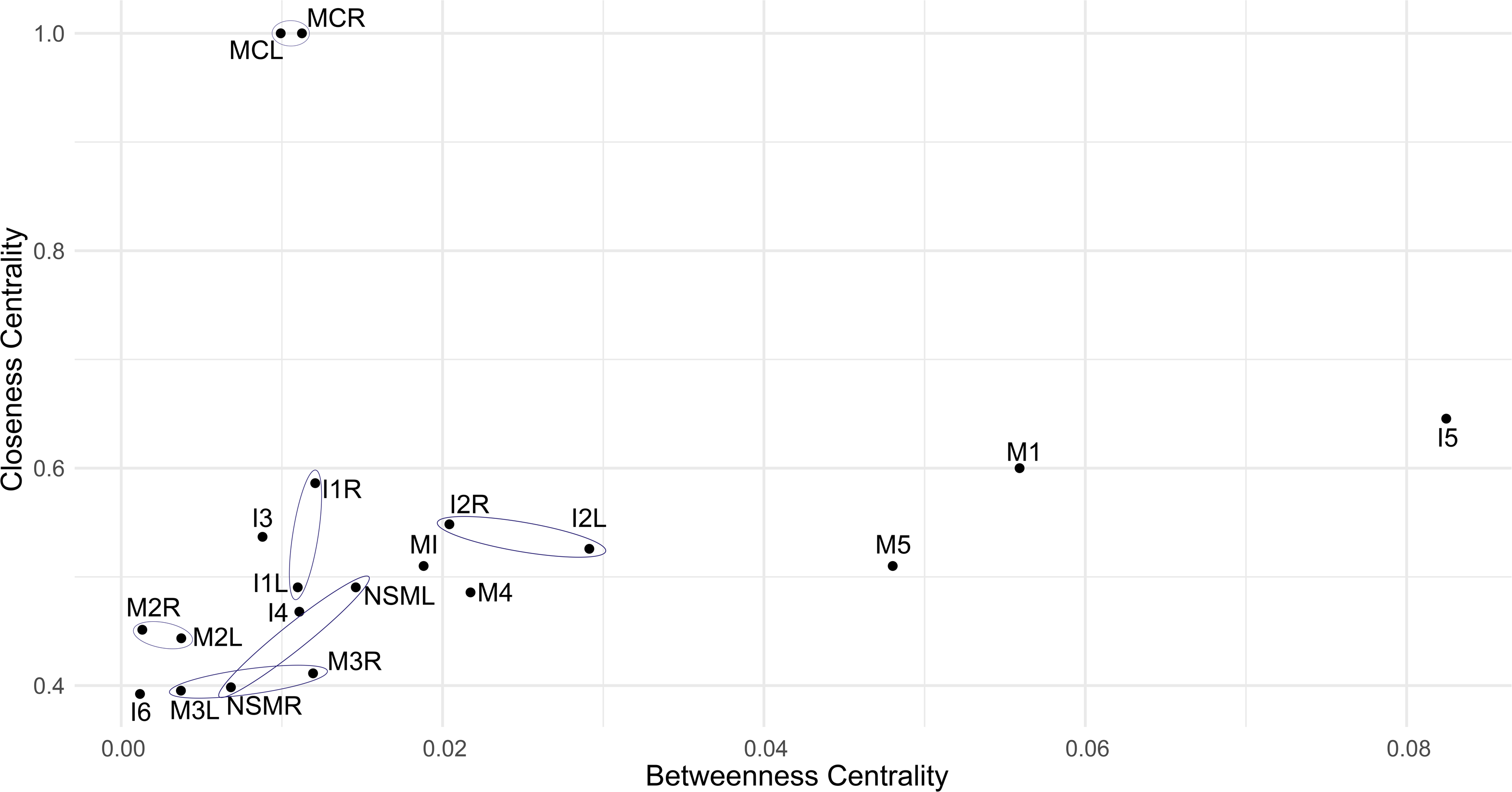
Betweenness centrality vs. closeness centrality for all C. elegans pharyngeal neurons. Plot showing the betweenness centrality (x-axis) vs. closeness centrality(y-axis) for each pharyngeal neuron. Blue ovals connect left/right homologous neuron pairs.

### Molecular basis for chemical and electrical communication in the pharyngeal connectome

As a basis for future molecular dissection of information flow within and from and to the pharyngeal nervous system, we extracted known gene expression patterns from wormbase.org, as they relate to neuronal communication, and overlaid them onto the pharyngeal connectome. The expression pattern analysis of genes encoding for neurotransmitter synthesizing enzymes, transporters, and receptors puts us in the unique position to assign neurotransmitter signaling pathways in the pharynx (Figure 10A)(**Supplemental data 6**)^34–36 37 38^. We found that 82/83 (98.8%) of chemical synapses have a matching expression pattern of a neurotransmitter with a cognate neurotransmitter receptor. Only a single synapse, NSM->I5, did not have matching neurotransmitter machinery. We found that the metabotropic receptors alone cover almost the entire pharyngeal nervous system, with 83% of chemical synapses having a matching expression pattern of NT with NTR (e.g. Ach + GAR). If we further limit our analysis to only larger connections, 100% of chemical connections >9 sections in weight have a matching neurotransmitter + neurotransmitter receptor pair. Of the 79 cholinergic and glutamatergic somatic neuronal classes, the metabotropic neurotransmission fraction is 73%.

**Fig. 10.**
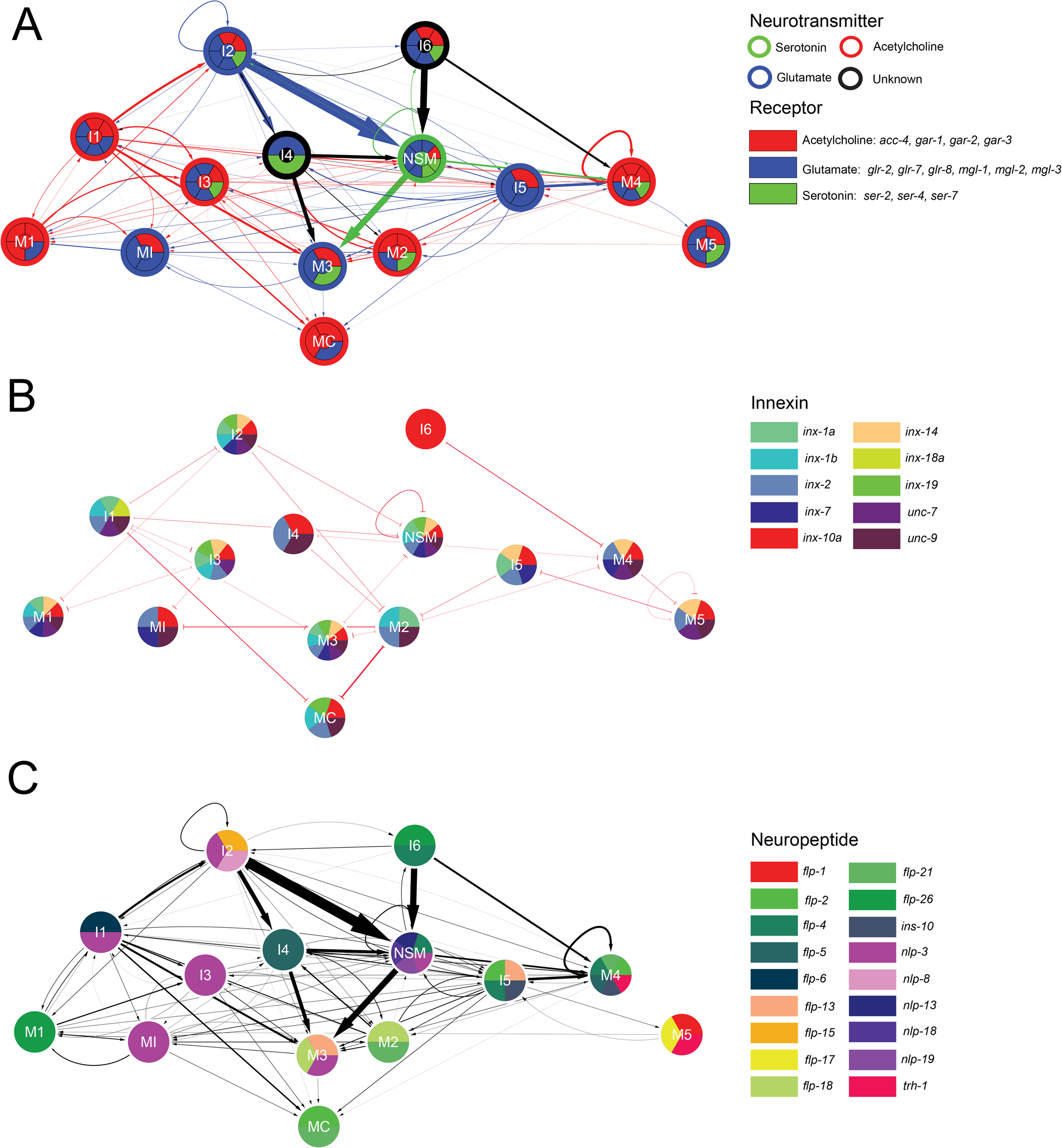
Molecular modes of communication within the pharynx. (**A**) Neurotransmitter identity (node outlines) and neurotransmitter receptor expression (pie graph sections) are shown for Acetylcholine, Glutamate, Serotonin. Edges are colored by the presynaptic neurotransmitter identity, legend to right. (**B**) Innexin expression patterns for pharyngeal neurons with gap junction connections between neurons (red), legend to right. (**C**) Neuropeptide expression patterns with chemical synapses between neurons (black lines and arrows), legend to the right.

As a first step to “de-orphanize” the pathway of electrical communication within the pharynx we made use of a recently published expression map of all innexin proteins, which are the building blocks of electrical synapses^39^. Each individual pharyngeal neuron expresses a unique set of innexin proteins, thereby providing each neuron with unique capacities for electrical synapse formation (Figure 10B).

The expression patterns of many neuropeptides have also been previously described^40^. Superimposed onto the pharyngeal connectome, these expression patterns reveal not only wide-spread usage of neuropeptides, but also demonstrates that, similar to innexins, each pharyngeal neuron expresses a unique combination of neuropeptides with the exceptions of MI and I3 (Figure 10C). It is important to appreciate that, based on a published precedent^41^, neuropeptide signaling will not be entirely restricted to within the pharynx, but neuropeptides may constitute a major means by which the pharyngeal nervous system communicates with the somatic nervous system.

## DISCUSSION

Our analysis of the pharyngeal nervous system connectome, a self-contained, autonomously acting unit within the *C. elegans* nervous system, reveals numerous novel insights into the function of individual neuron types and demonstrates themes of neuronal circuit organization, inviting speculation about neuronal circuit development and evolution.

We discovered that most neurons in the pharyngeal nervous system have likely sensory function and that all of them directly innervate end organs. These insights substantially extend the previous analysis of the anatomy of the pharyngeal connectome^6^. Moreover, our network analysis reveals that individual neurons are topologically organized into functional modules that innervate distinct functional domains of the pharynx (corpus, isthmus, terminal bulb). Sensory stimuli can therefore be quickly transduced to motor output via shallow processing depth within each module. The mechanism(s) for signal transduction by the primitive sensors in this tissue remain to be explored. Given the lack of a true cilium or basal body, it remains unclear whether any of the BBS molecular machinery could play a role here^25^. While the overall network structure is shallow, there is also extensive cross-connectivity between neurons and network analysis reveals a small number of neurons that appear to be central to overall network structure. Previous cell ablation studies yielded weak or no pumping phenotype for most neurons and strong phenotypes for only a subset; technical limitations or compensatory changes may have obscured the function of others. Acute silencing of these neurons (e.g. I5 or M5) may be the best way to probe our prediction about their functional relevance for pharyngeal behavior.

Essentially all synaptic contacts between distinct neuron classes in the *C. elegans* nervous system are made *en passant.* As originally pointed out by John White^42^, there are two critical choice points for the establishment of such *en passant* synaptic circuitry in the somatic nervous system. First, a neuron makes a selective fasciculation choice to pick the neighborhood where its process will be placed, thereby restricting synaptic targeting choice to a select number of neurons. Second, within the chosen neighborhood, neurons in the somatic nervous system then make synaptic contacts with about one fifth of their neighbors^42^. We found that pharyngeal neurons more frequently come into contact with their nearby neighbors and more frequently form synapses with physically adjacent neurons. Hence, pharyngeal neurons appear to be endowed with less synaptic targeting specificity as neurons in the somatic nervous system. This observation corroborates the critical importance of proper neighborhood choice, i.e. selective fasciculation during process outgrowth, in synaptic target choice. It will be fascinating to see whether selective process fasciculation involves a selective sorting of jointly growing axons or whether fascicles are built sequentially, such that a temporally controlled outgrowth and access to preexistent bundles determines neuronal process adjacencies.

Both the sensory/inter/motorneurons features of most pharyngeal neurons, as well as the apparently relative promiscuity in target selection, offer a novel perspective on the pharyngeal nervous system that relates to current thoughts on the evolution of multicellularity and the evolution of nervous systems. The first cell types to have specialized from each other are generally thought to be primitive neurons that specialized in their ability to perceive and relay signals to contractile myoepithelial cells, the second specialized cell type thought to have come into existence^43^. Such “ur-“neurons were therefore simultaneously serving as sensory and motor neurons that may also have formed simple nerve nets to relay information among each other, making these ur-neurons also primitive interneurons. Seen from this perspective, the architecture of the pharyngeal nervous system, composed of such primitive, sensory/inter/motorneurons which innervate myoepithelial target cells (such as the pharyngeal muscle) is reminiscent of the hypothetical architecture of a primitive, ancestral nervous system.

## MATERIALS AND METHODS

### Electron micrograph reconstruction

Three series of electron micrograph prints were used to reconstruct the entire adult hermaphrodite pharyngeal nervous system of *C. elegans* (Figure 2A)^6^. These EM samples were prepared as previously described^44^. The N2T series covers the procorpus and the N2W series covers the metacorpus, isthmus, and posterior bulb. The JSA and N2W series both cover the pharyngeal nerve ring, a complex commissure of axonal processes in the pharyngeal metacorpus, is the most complex neuropil of the pharyngeal nervous system. We averaged the connectivity of two nerve ring reconstructions, and then combined these connectivity data with the remainder of the N2W and N2T series to generate a complete pharyngeal connectome. Connections present in only the JSA or N2W series are represented as 0.5*edge weight (**Supplemental data 3**). Digitized electron micrographs were aligned using TrakEM2^45^, and reconstructed using Elegance^46^. Skeleton diagrams of neurons, chemical synaptic and gap junction connectivity between cells, and weighted adjacency matrices were generated and are available at www.wormwiring.org. Volumetric tracings of electron micrographs were performed using TrakEM2^45^. Images of electron micrographs used in this reconstruction can be accessed at www.wormimage.org.

### Connectivity and adjacency analysis

The connectivity data that correspond to this manuscript can be found at www.wormwiring.org and in the supplemental data. Chemical synapses and gap junctions were annotated using previously established ultrastructural criteria determined in *C. elegans*^3^. The anatomical strength of connection was estimated by summing the number of serial section electron micrographs where presynaptic specializations or gap junctions were observed. Chemical synaptic and gap junction connectivity were treated as weighted directed and weighted undirected graphs, respectively. Nodes in the graph include neuron, muscle, epithelial, marginal, and gland cells. Volumetric reconstructions were completed by manually tracing cell membranes using TrakEM2^45^. Adjacencies of neurons were determined by applying a modified python script to extract cell-cell contacts from TrakEM2 reconstructions^30^. Graph-theoretic analyses were performed using a combination of python and MATLAB scripts as well as the network analyzer plugin for Cytoscape^47^. Statistical analyses were completed using a combination of Python, MATLAB, and R^48^; figures were generated using Python and ggplot2^49^.

### Strains and transgenes

Worms were maintained using standard methods^50^. Wild-type Bristol N2 animals were grown at 20 °C on *Escherichia coli (OP50)*-seeded nematode growth medium plates as a food source. Each transgenic stain was generated by microinjecting^51^ 33 ng/ul of cytoplasmic marker (e.g. *pdfr-1::tagRFP)*, 7 ng/ul synaptic marker (e.g. *gur-3::GFP::CLA-1),* 50 ng/ul of pRF4 (*rol-6* co-injection marker), and 10 ng/ul of carrier DNA (pBluescript) for a total concentration of 100 ng/ul. A list of strains used in this study are listed in **Supplemental data 9**. To generate cytoplasmic markers, promoter fragments were amplified from N2 genomic DNA and cloned into 5’-*SphI*, 3’-*XmaI* digested vectors expressing *tagBFP* or *tagRFP* using Gibson assembly or by T4 ligation using 5’-*SphI*, 3’-*XmaI* digested promoter fragments. To generate synaptic reporters, promoter fragments were amplified from N2 genomic DNA and cloned into 5’-*SphI*, 3’-*XmaI* digested vectors expressing *mCherry::RAB-3*, *GFP::RAB-3*, or *GFP::CLA-1* by T4 ligation using 5’-*SphI*, 3’-*XmaI* digested promoter fragments. A list of plasmids used in this study are listed in **Supplemental data 10**.

### Light microscopy

Young-adult worms were anaesthetized using 100 mM Sodium azide (NaN_3_) and mounted onto glass slides with 5% agarose pads and visualized using fluorescence microscopy with Nomarski optics. Strains expressing *RAB-3::mCherry* were imaged using a Zeiss Axioimager Z1 with Apotome. All other strains were imaged using a Zeiss 880 laser-scanning confocal microscope. To analyze multidimensional data, we generated maximum-intensity projections using ImageJ and quantified distinct dots (corresponding to synaptic puncta). Adobe Illustrator CC was used to create figures.

## Acknowledgements

We thank John G. White and Jonathan Hodgkin for their help in transferring archival TEM data from the MRC/LMB to the Hall Lab at the Albert Einstein College of Medicine for long-term curation and study. Christopher Brittin was influential in designing and creating www.wormwiring.org and providing software support. Christopher Crocker provided artistic assistance, with some figures adapted from original www.wormatlas.org drawings. We thank Chi Chen technical injection assistance. We also thank Hannes Bülow, Emily Bayer, Berta Vidal-Iglesias, and Raphael Cohn for comments and discussion regarding this manuscript. This work was supported by the G. Harold and Leila Y. Mathers Charitable Foundation (S.W.E.), R24OD010943 (D.H.H.), a T32GM007491 award to the Albert Einstein College of Medicine (S.J.C.), and a postdoctoral fellowship (5F32MH115438) to S.J.C. O.H. is an Investigator of the Howard Hughes Medical Institute.

